## Supporting Information for "Multivalent Interactions between Molecular Components Involved in Fast Endophilin Mediated Endocytosis Drive Protein Phase Separation"

**Supplementary figures**

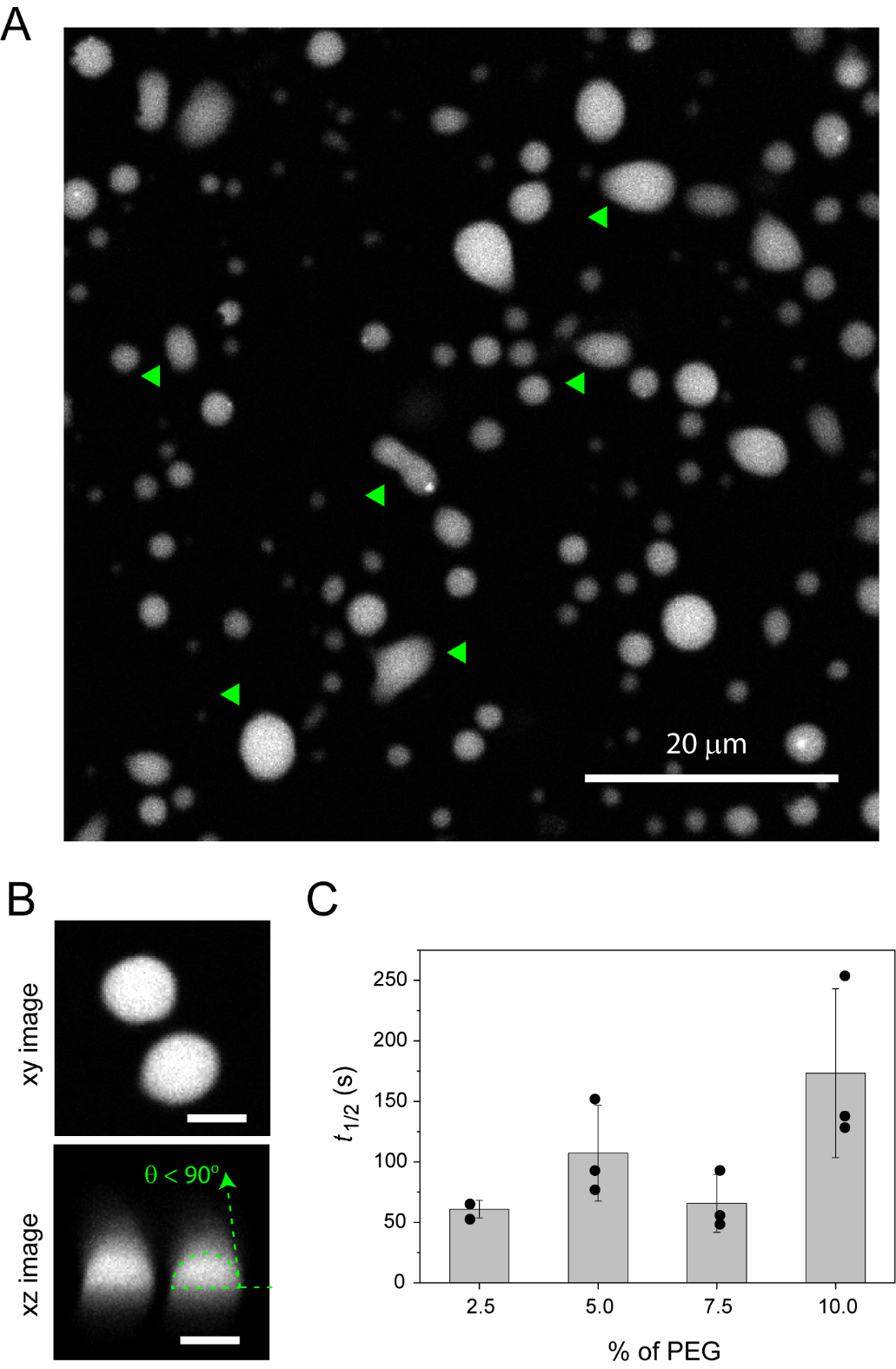

**Supplementary Figure 1:** Endophilin droplets exhibit liquid-like properties. **A.** Confocal image showing coalescence behavior of the endophilin droplets on the glass surface. Green arrows indicate adjacent droplets coalesced to form large, non-circular shapes. **B.** Wetting behavior of the endophilin droplets on glass surface. xy and xz images showing spherical shape of the droplets that form a contact angle θ < 90⁰ on the glass surface. **C.** Time constants (*t*_1/2_) obtained from exponential fittings of FRAP recovery profiles for endophilin droplets formed at various PEG concentrations. Bar graph represents mean and standard deviations from three independent FRAP experiments.

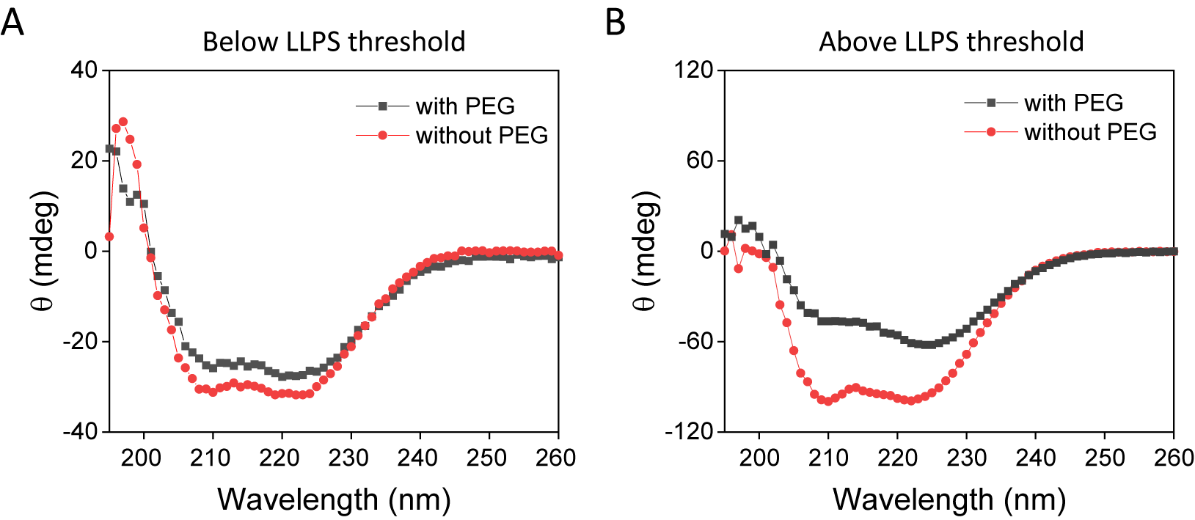

**Supplementary Figure 2:** Circular dichroism spectra of endophilin in the presence and absence of 10% PEG. Spectra were recorded at a protein concentration of 5 µM (A) which is below the phase boundary and 20 µM (B) which is above the phase boundary.

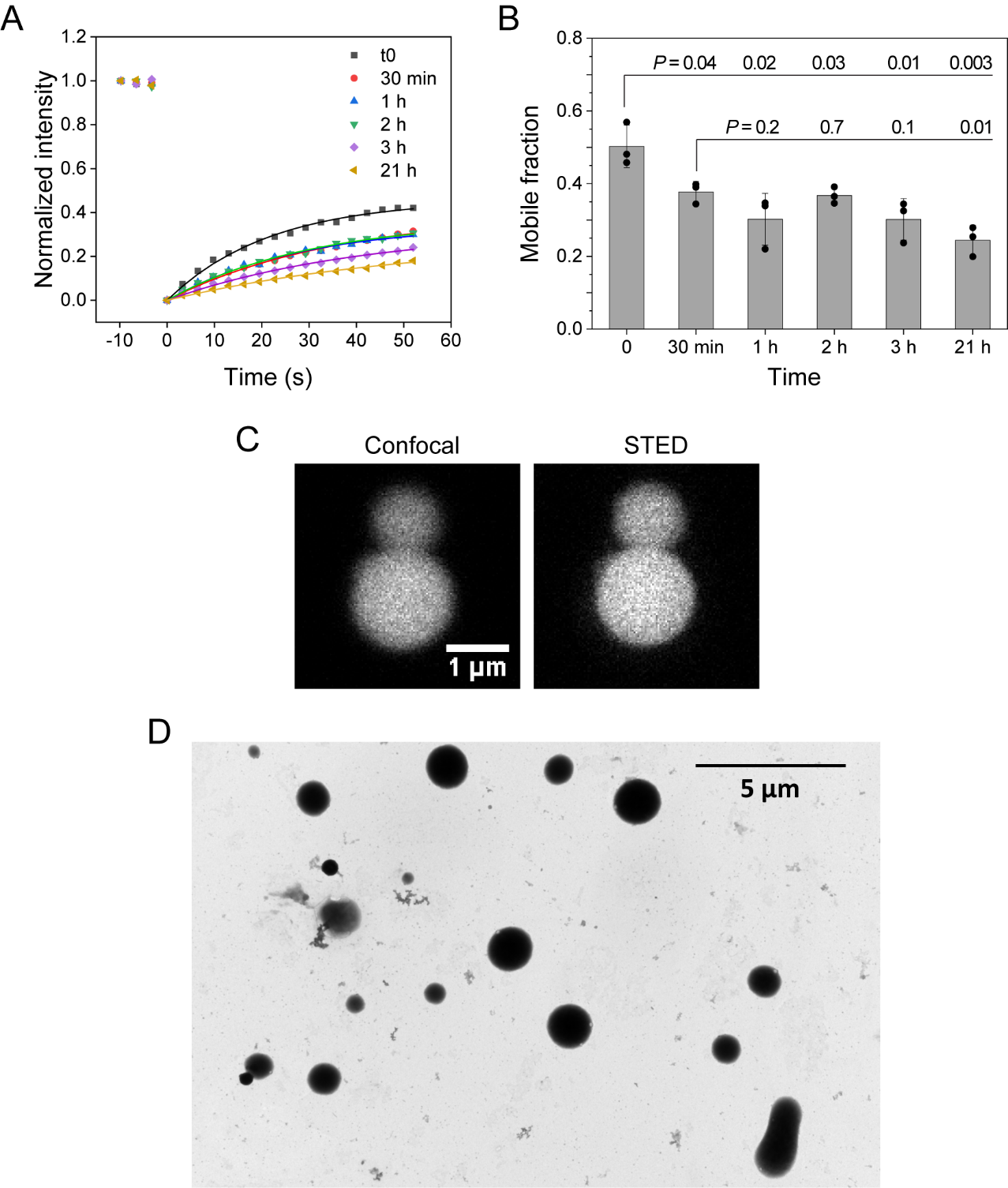

**Supplementary Figure 3**: Material property characterized for endophilin droplets by optical and electron microscopy show homogeneous, liquid-like droplets. **A.** Fluidity of the endophilin droplets formed in the presence of 5% PEG monitored over the course of 21 h. FRAP recovery profiles of experiments performed at different time points after introducing PEG to the protein solution. **B.** Mobile fractions determined from fitting of the FRAP data with single exponential model. An average of three independent FRAP experiments has been plotted along with the data points, error bars represent standard deviations. The *P* values (two-tailed) are obtained from Student’s *t*-test. **C.** Confocal (left) and STED (left) microscopy images of endophilin droplets (25 µM, 10% PEG) containing Alexa 488-labeled endophilin (1 µM). **D.** Negative stain TEM image of endophilin LLPS condensates formed in the presence of 10% PEG does not show any heterogeneity.

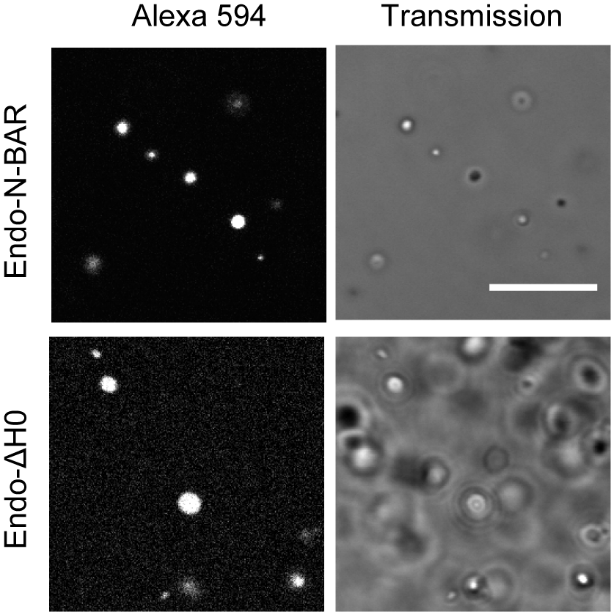

**Supplementary Figure 4:** Droplets formed by the truncation mutants of endophilin in the presence of 10% PEG. Left, confocal images of the droplets recorded in the presence of Alexa 594-labeled proteins (4 mol %), keeping total protein concentration of 25 µM. Right, corresponding transmission images. Scale bar 10 µm.

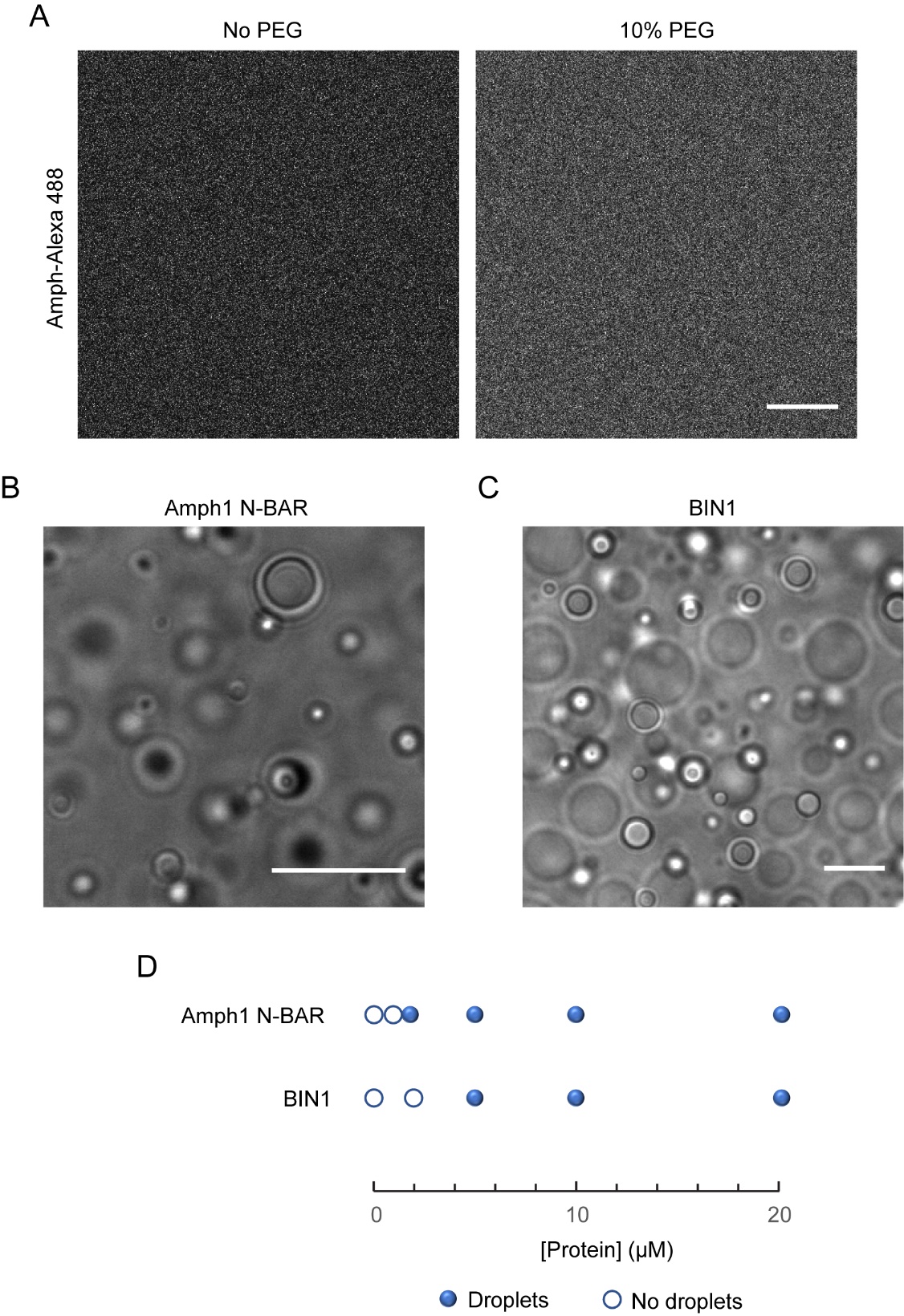

**Supplementary Figure 5**: Full length amphiphysin (Amph1) does not form liquid-like droplets whereas the N-BAR domain of Amph1 and full-length BIN1 (isoform 9) undergoes LLPS in the presence of PEG. **A.** Confocal images of solutions containing amphiphysin (25 µM total protein concentration; 96% unlabeled, 4% Alexa 488 labeled) in the presence and absence of PEG (10 % w/v). **B** and **C.** Transmission images of droplets formed by Amph1 N-BAR (25 µM) and full length BIN1 (35 µM) respectively, in the presence of 10% PEG. **D.** LLPS boundary of Amph1 N-BAR and BIN1 in the presence of 10% PEG. All scale bars are 10 µm.

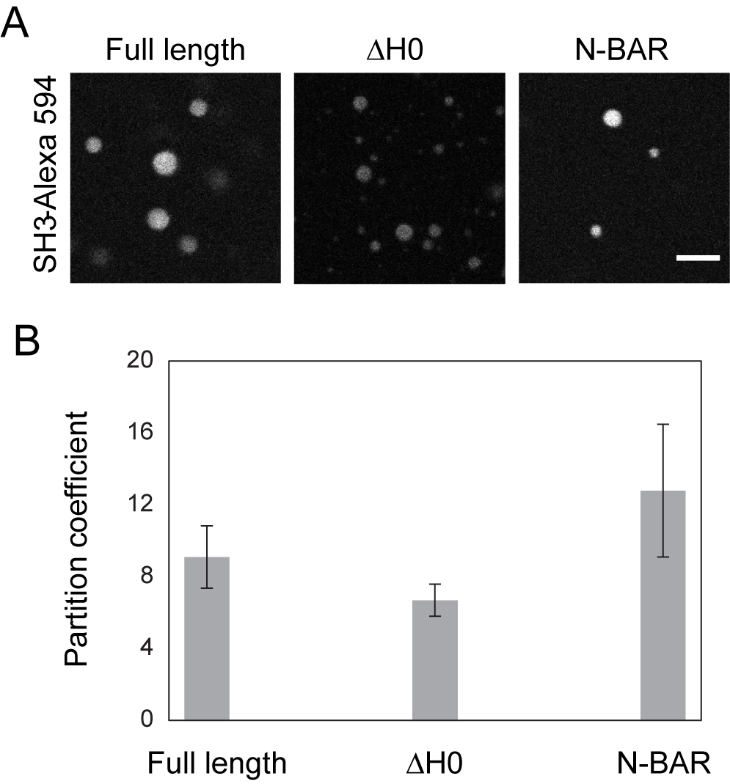

**Supplementary Figure 6:** Partitioning of SH3 domain of endophilin into droplets formed by full length endophilin and two truncation mutants. **A.** Confocal images of the droplets formed by 25 µM of the proteins (10% PEG) in the presence of Alexa 594-labeled SH3 domain (1 µM). Scale bar 5 µm. **B.** Bar plot showing apparent partition coefficients determined from Alexa 594 intensities inside and outside droplets. Data is presented as mean ± standard deviations.

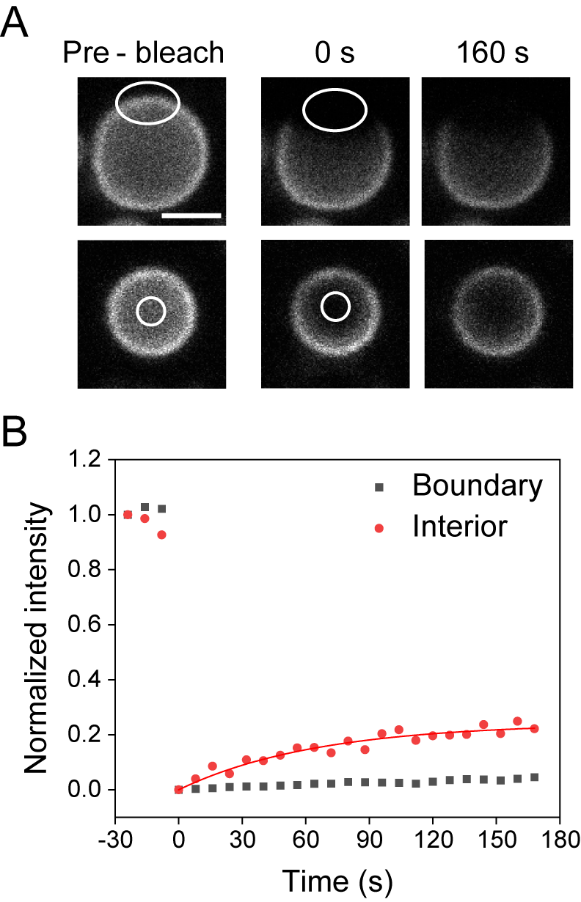

**Supplementary Figure 7**: FRAP studies on anisotropically partitioned Alexa 488-labeled amphiphysin (200 nM) into endophilin droplets (25 µM, 10% PEG). The boundary and the interior of the droplets were separately photobleached (A) and their FRAP profiles were plotted (B). Scale bar 2.5 µm.

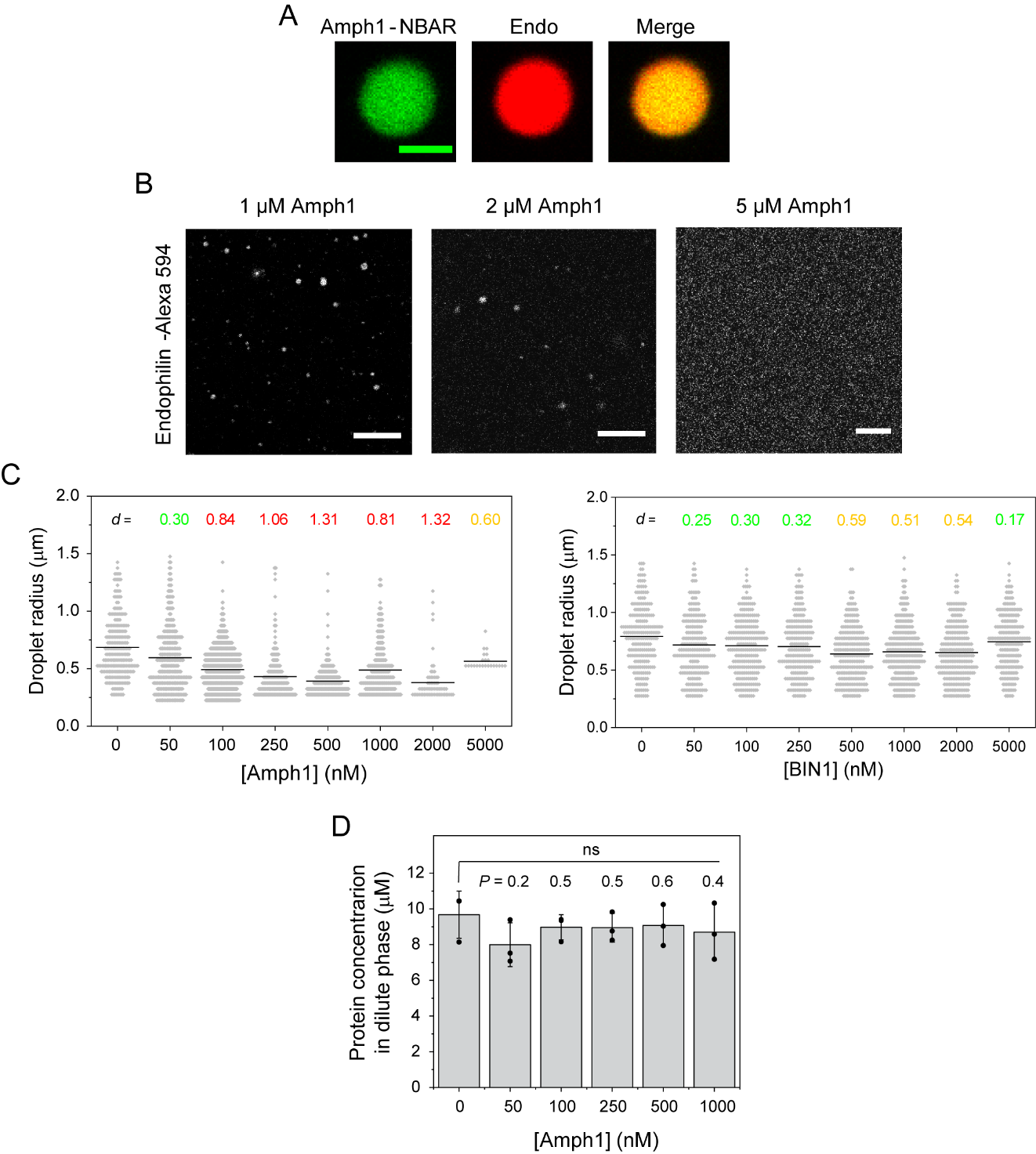

**Supplementary Figure 8: A.** Distribution of N-BAR domain of amphiphysin (Alexa 488 labeled, 200 nM) within endophilin droplets (25 µM, along with 4% Alexa 594 labeled) formed in the presence of 10% PEG. Scale bar 2.5 µm. **B.** Confocal images of showing inhibition of endophilin droplet formation in the presence of amphiphysin. Very few droplets were observed in the presence of 1 and 2 µM of amphiphysin whereas no droplets formed in the presence of 5 µM amphiphysin. Scale bars 10 µm. **C.** Size distribution of endophilin droplets in the presence of various concentrations of Amph1 (left) and BIN1 (right). The numbers denote Cohen’s *d* values that represent the ‘effect size’ and are colored to show whether effect is small (*d*~0.2, green), medium (*d*≥ 0.5, yellow), or large (*d*≥ 0.8, red) according to Cohen’s convention. **D.** Dilute phase protein concentration estimated for endophilin-PEG LLPS system in the presence of various concentrations of Amph1. *P* values (two-tailed) obtained from Student’s *t*-test indicate no significant change in the dilute phase concentration takes place in the presence of Amph1.

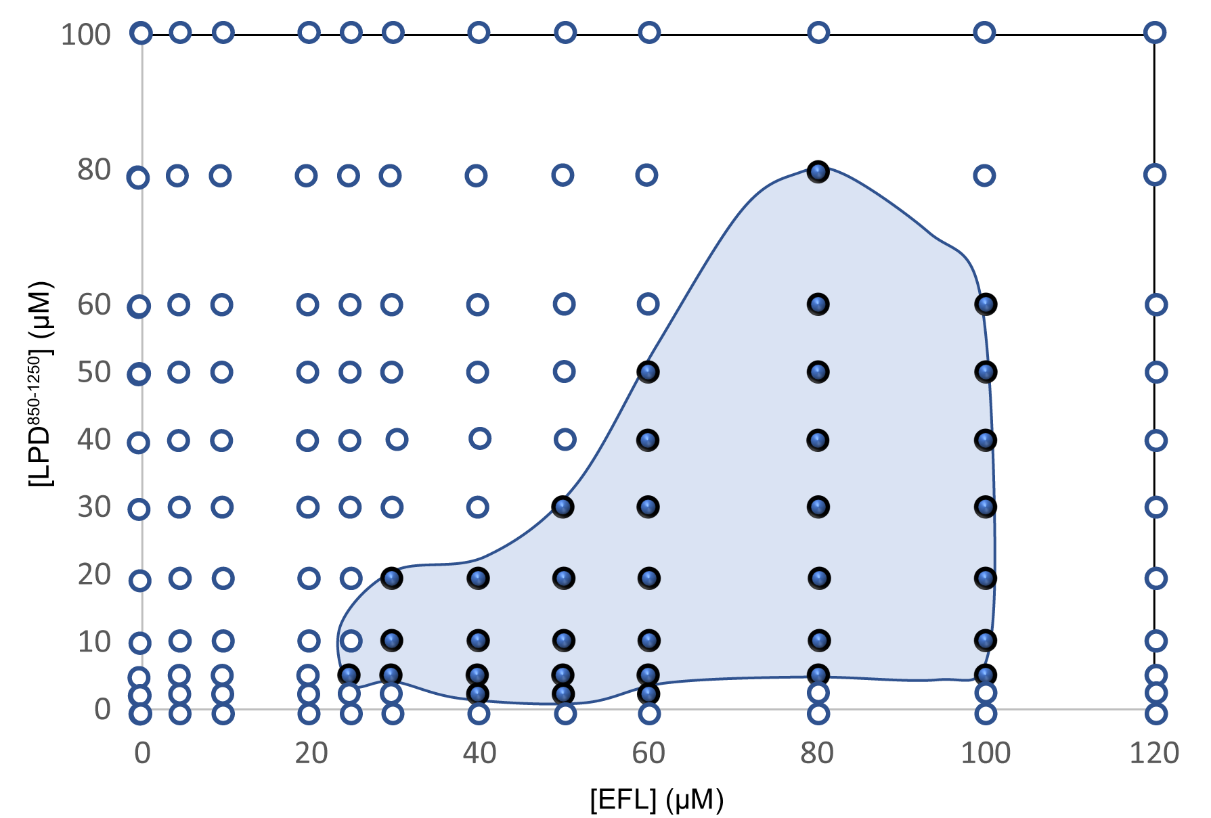

A

B

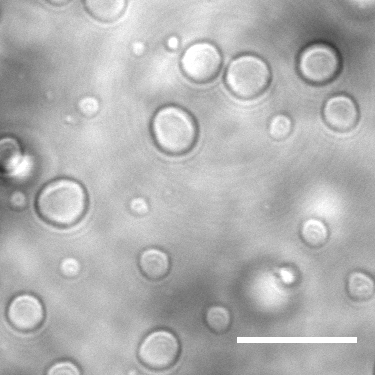

**Supplementary Figure 9: A.** Complete phase diagram for the endophilin/LPD^850-1250^ LLPS system that shows closed loop phase behavior. Filled circles represent observation of condensates whereas empty circles represent no phase separation. The enclosed area roughly indicates the boundary of the concentration regime that shows LLPS. **B.** Transmission images of LLPS droplets formed by VASP (20 µM) and LPD^850-1250^ (40 µM) upon mixing. Scale bar 10 µm. All experiments were performed in 20 mM HEPES buffer, 150 mM NaCl, 1 mM TCEP, pH 7.4 and at room temperature.

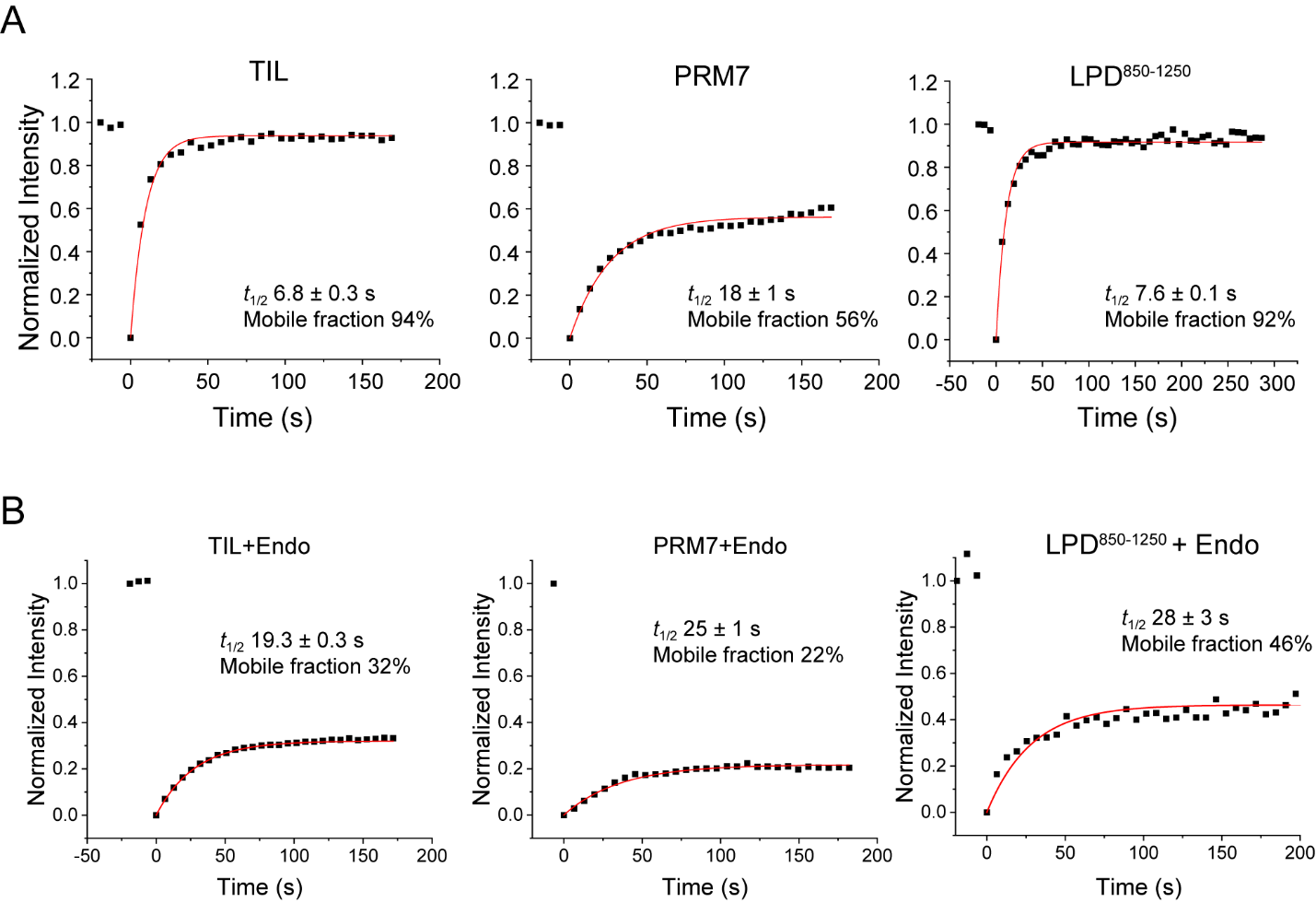

**Supplementary Figure 10:** FRAP profiles and their single-exponential fits for TIL (Alexa 488 labeled), PRM7 (Alexa 633 labeled), and LPD^850-1250^ (Alexa 647-labeled) tethered onto solid supported bilayers via His_6_ tags in the absence (A) and presence of endophilin ΔH0 (2.5 µM) (B).

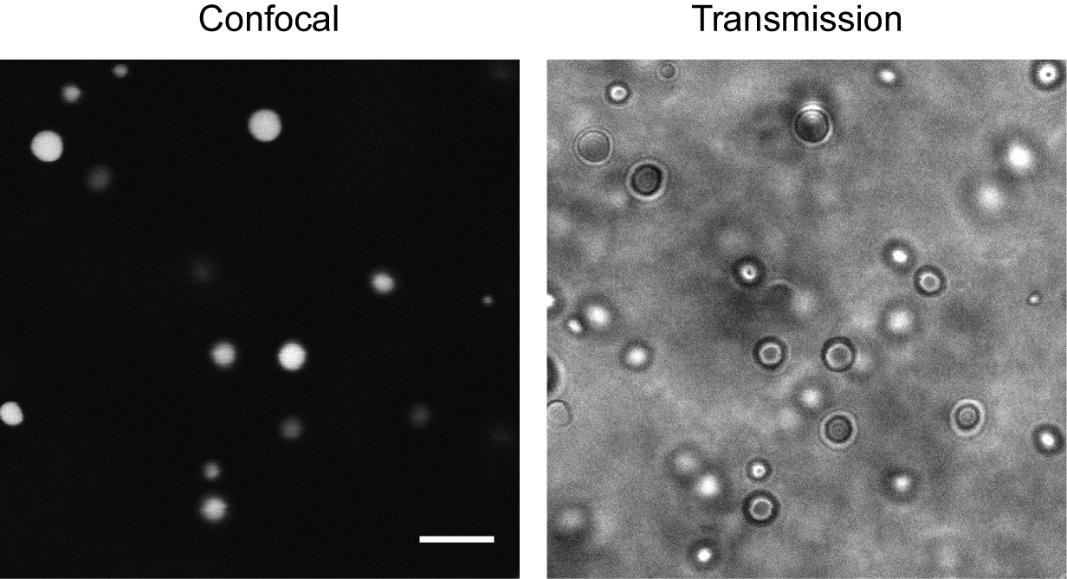

**Supplementary Figure S11**: Confocal fluorescence image (left) and corresponding transmission image (right) of liquid-like droplets formed by endophilin ΔH0 and TIL (Alexa 488 labeled) when mixed at equimolar concentrations (75 µM). Scale bar 10 µm.

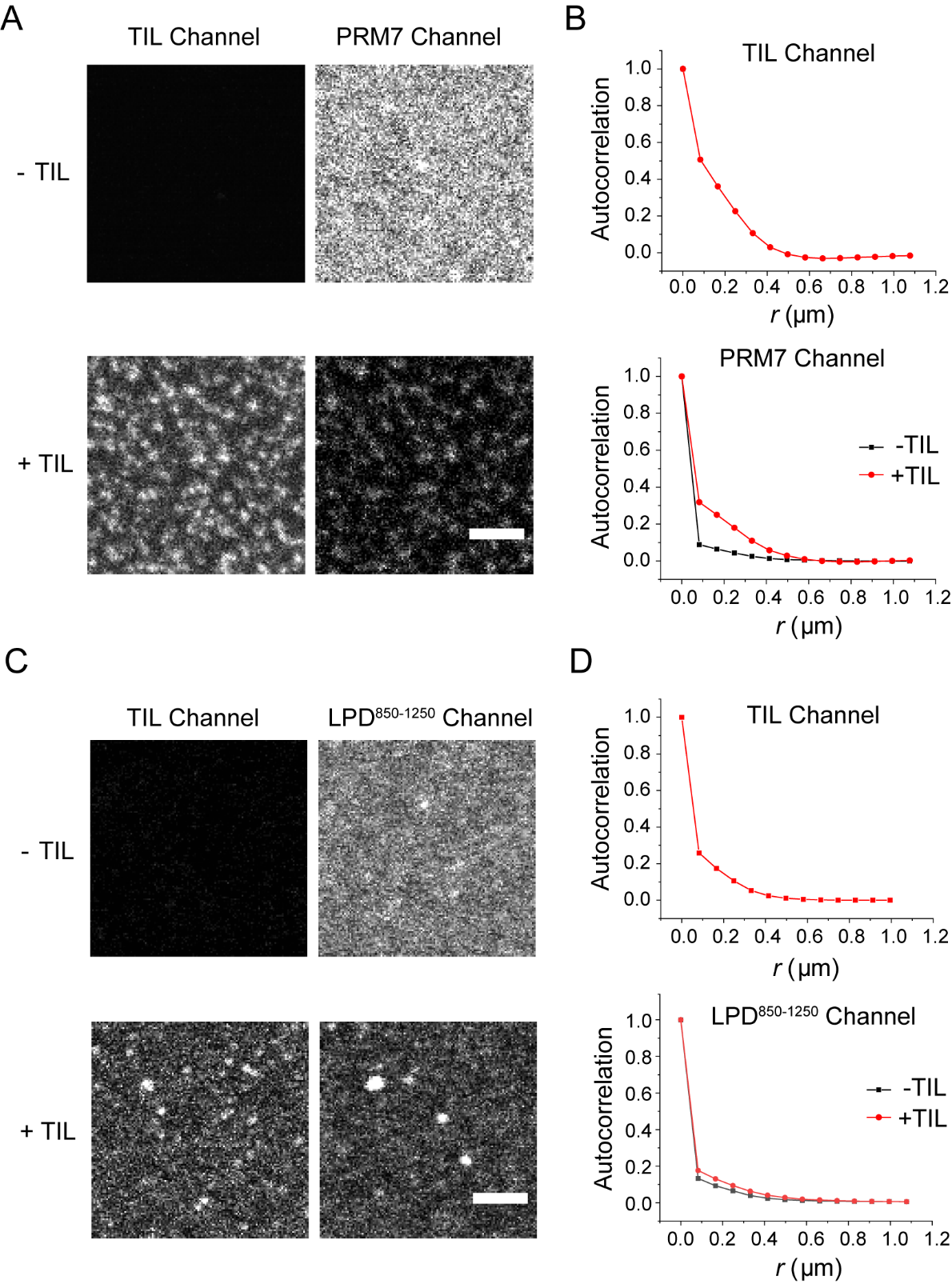

**Supplementary Figure 12**: Clustering of TIL and LPD on the bilayer in the absence of endophilin. **A.** Confocal images of bilayers with conjugated PRM7 (500 nM) before (top) and after (bottom) addition of TIL (500 nM). Scale bar 2.5 µm. **B.** Radial distribution autocorrelation functions showing the extent of clustering in the TIL channel (top) and PRM7 channel (bottom) before and after addition of TIL. **C.** Bilayers with conjugated LPD^850-1250^ (250 nM) before (top) and after (bottom) addition of TIL (50 nM). Scale bar 2.5 µm. **D.** Autocorrelation functions in the TIL channel (top) and LPD^850-1250^ channel (bottom) before and after addition of TIL.

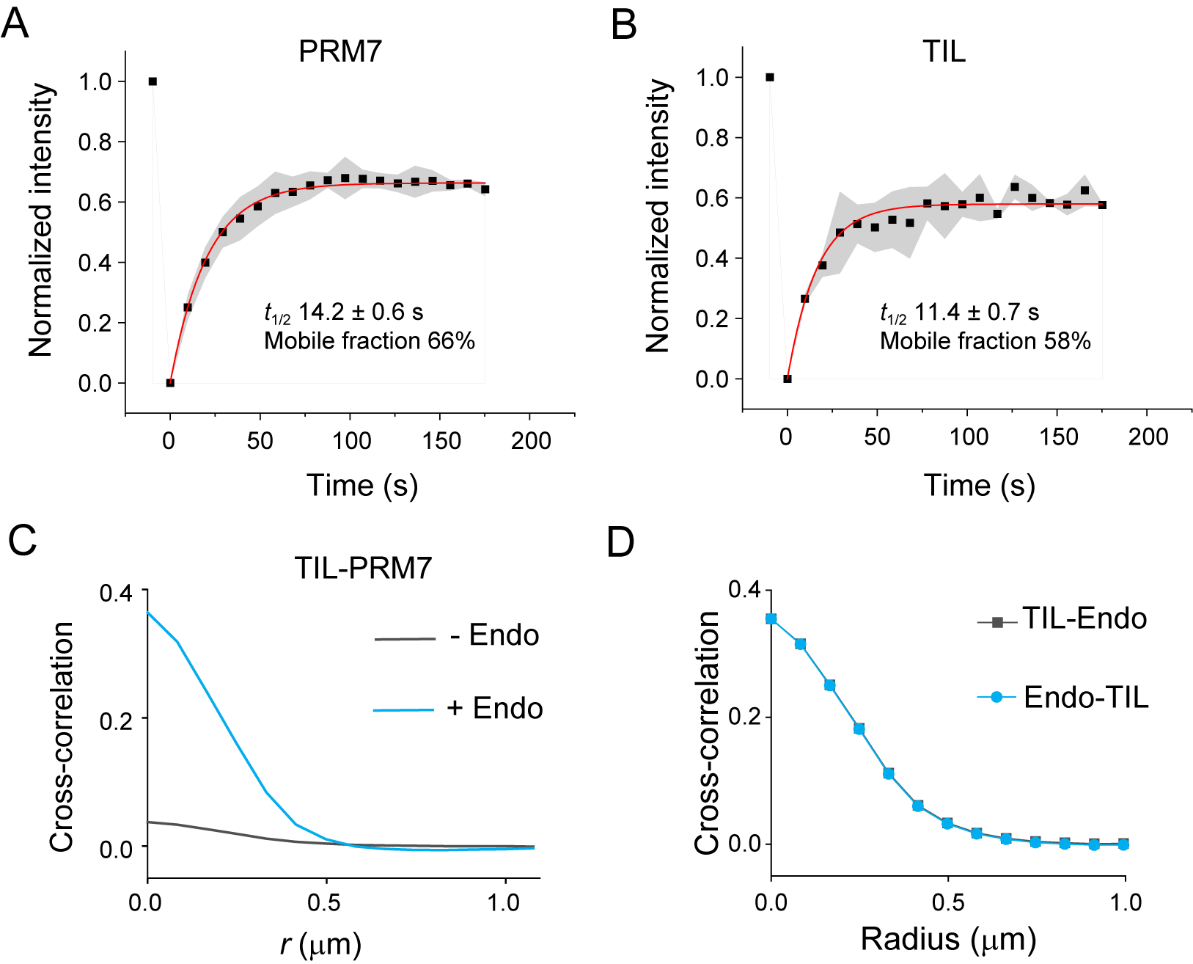

**Supplementary Figure 13**: FRAP profiles for solid supported bilayers with conjugated TIL-Alexa 488 and PRM7-Alexa 633. Both PRM7 (A) and TIL (B) shows photorecovery indicating that they are mobile on the bilayer. C. Cross-correlation plot showing the extent of co-localization between TIL and PRM7 channels increases after addition of endophilin. D. Cross-correlation functions for endophilin-TIL and TIL-endophilin are plotted together to show that they overlap with each other.

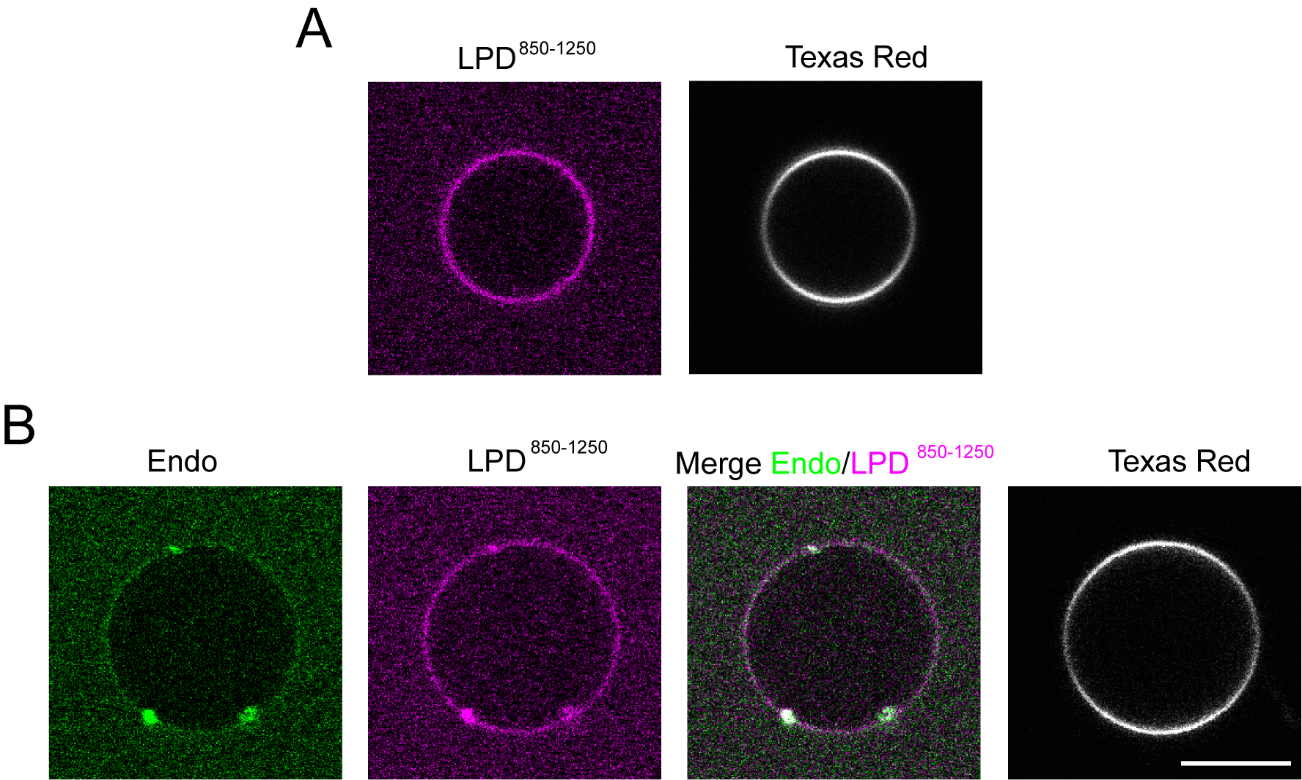

**Supplementary Figure 14**: Endophilin and LPD850-1250 undergo clustering on GUVs. Confocal images of GUVs (DOPC/NTA/Texas red in 98.8:1:0.2 molar ratio) with conjugated LPD^850-1250^-His_6_ (Alexa 647) in the absence (A) and presence (B) of endophilin-Alexa 488 (200 nM). Scale bar 10 µm. Clustering of both endophilin and LPD^850-1250^ occurs on GUVs in the presence of endophilin whereas no changes in the membrane structure (no interior nor exterior tubulation or budding) were observed in the lipid (Texas Red) channel.

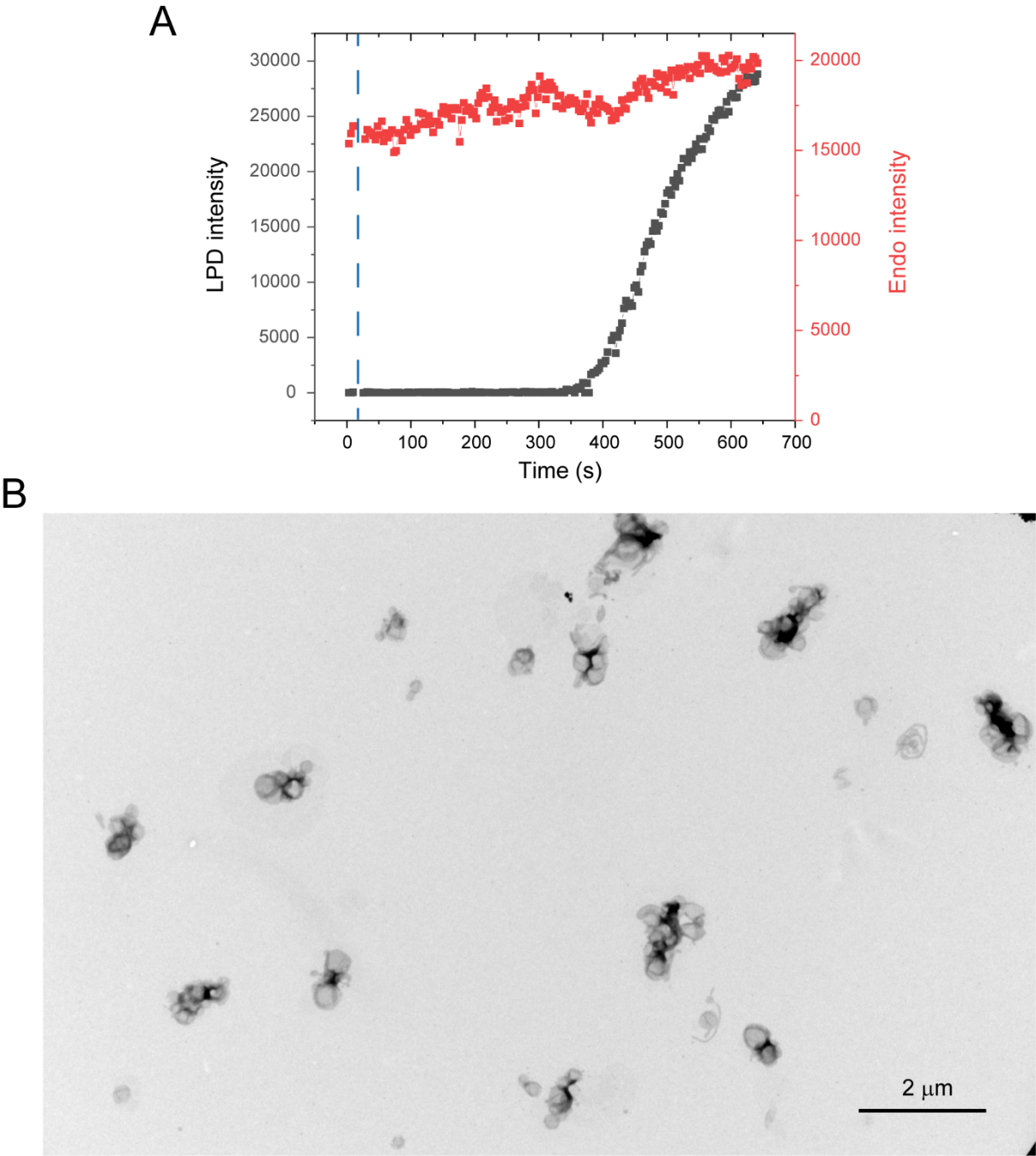

**Supplementary Figure 15**: **A.** Intensity profile of LPD^850-1250^ (Alexa 647) (grey) and endophilin (Alexa 488) (red) channels on the GUV surface indicating that addition of LPD does not cause a change in the membrane density of endophilin. The blue dotted line indicates the time point of LPD addition. **B.** TEM image of clusters of liposomes caused by membrane adherence in the presence of endophilin and LPD^850-1250^.

**Supplementary Tables**

**Supplementary Table 1:** Apparent partition coefficients (***K*_app_**) for client proteins/peptides into endophilin droplets.

|  | ***K*_app_** |
| --- | --- |
| **PRM7** | 26 ± 4 |
| **TIL** | 138 ± 40 |
| **LPD850-1250** | 515 ± 60 |
| **BIN1** | 267 ± 20 |
| **Amph** | 352 ± 80 |
| **FBP17** | 1128 ± 350 |
| **BSA** | 2.9 ± 0.4 |
| **Syn** | 2.67 ± 0.02 |

**Supplementary Table 2:** Correlation length (*r*) obtained from single exponential fits of the auto-correlation functions to estimate the extent of endophilin induced clustering of TIL, PRM7, and LPD^850-1250^ on lipid bilayers (Figure 4F).

|  | *r* |
| --- | --- |
| **TIL + 0 µM Endo** | 0.029 ± 0.002 |
| **TIL+ 1 µM Endo** | 0.089 ± 0.006 |
| **TIL+ 2.5 µM Endo** | 0.142 ± 0.007 |
| **PRM7 + 0 µM Endo** | 0.036 ± 0.003 |
| **PRM7 + 1 µM Endo** | 0.119 ± 0.01 |
| **PRM7 + 2.5 µM Endo** | 0.15 ± 0.01 |
| **LPD850-1250 + 0 µM Endo** | 0.035 ± 0.003 |
| **LPD850-1250 + 1 µM Endo** | 0.118 ± 0.009 |
| **LPD850-1250 + 2.5 µM Endo** | 0.181 ± 0.008 |

**Supplementary Table 3:** Correlation lengths (*r*) for TIL and PRM7 co-coupled on bilayers in the presence and absence of endophilin (Figure 5C).

|  | - **Endo** | **+ Endo** |
| --- | --- | --- |
| **TIL** | 0.057 ± 0.001 | 0.154 ± 0.001 |
| **PRM7** | 0.028 ± 0.001 | 0.069 ± 0.001 |
| **Endo** | - | 0.153 ± 0.001 |

**Supplementary Table 4:** Correlation lengths (*r*) in the presence and absence of TIL on bilayers containing pre-existing clusters of either endophilin/PRM7 (Figure 5F) or endophilin/LPD^850-1250^ (Figure 5I).

|  | **- TIL** | **+ TIL** |
| --- | --- | --- |
| **TIL** | - | 0.139 ± 0.001 |
| **PRM7** | 0.073 ± 0.001 | 0.177 ± 0.002 |
| **Endophilin** | 0.115 ± 0.001 | 0.219 ± 0.003 |
|  | **- TIL** | **+ TIL** |
| **TIL** | - | 0.111 ± 0.001 |
| **LPD^850-1250^** | 0.11851 ± 0.00139 | 0.217 ± 0.001 |
| **Endophilin** | 0.09261 ± 0.00126 | 0.164 ± 0.001 |

**Supplementary Methods**

**Purification of VASP**

Human VASP (wild type) was expressed in BL21(Codon plus DE3) cells as a N-terminal GST fusion protein. Cells were lysed by sonication in a buffer containing 300 mM NaCl, 50 mM Tris, 20 mM Imidazole, 2 mM DTT, 5 % Glycerol, pH 8, with 1 mM PMSF added. Soluble fraction of lysate was separated by centrifugation at 14,000 rpm for 1 hour, and the supernatant was filtered using 0.22 µm filter. The clarified lysate was incubated with Glutahione Sepharose beads (GE) overnight. The bound protein was washed with the lysis buffer and was incubated with PreScission protease overnight at 4 °C. The supernatant was confirmed via SDS-PAGE to contain cleaved VASP, and it was further refined by passing through a size exclusion column (Superdex 200), using 25 mM HEPES, 200 mM NaCl, 1 mM EDTA, 1 mM TCEP, 5% glycerol, pH 7.5.

**Stimulated-emission Depletion (STED) Microscopy**

STED images of the endophilin droplets were collected with a Leica CS SP8 STED 3X microscope (Leica Microsystems) using HC PL APO 100x/1.40 NA oil immersion objective (Leica). Alexa 488 labeled endophilin was excited at 490 nm using a tunable laser to collect both the confocal and STED images and emission was collected between the 505-560 nm wavelength range. STED depletion was achieved using a 592 nm continuous wave laser set at 40% laser power. Images were collected at 1016x1016 format using pixel dwell time of 600 ns. Droplet samples were formed by mixing endophilin (25 µM, containing 1 µM Alexa 488 labeled protein) and PEG (10% w/v) in HEPES buffer and the mixture was placed into a closed chamber created by sandwiching two coverslips (25 × 40 mm^2^, Fisher Scientific) using vacuum grease.

**Circular Dichroism (CD) Spectroscopy**

CD spectra were recorded with an AVIV CD spectrophotometer (Aviv Biomedical, NJ) using a High Precision Quartz Suprasil cuvette (Hellma Analytics) at 25 °C. Endophilin was mixed with phosphate buffer (10 mM sodium phosphates, 150 mM NaCl, pH 7.4) with or without containing 10% PEG (w/v) up to a final concentration of either 5 µM or 20 µM before recording CD spectra.

**Droplet size determination from confocal images**

Droplet size was determined from the confocal images using ImageJ. First, a binary image was created from an experimental image by applying Bandpass filter FFT method followed by threshold adjustment. Next, areas of all individual circular objects in the images were determined separately using the Analyze Particle tool. From the area, the diameter of each particle was determined. Due to large sample size (> 200 droplets in each group) we reported the actual effect size instead of *P* values obtained from a *t*-test, that resulted into extremely low *P* values (in the range of 10^-10^ to 10^-20^) in many cases. The effect size between two groups 1 and 2 was determined by calculating Cohen’s *d* values as follows ^1^,

$$d=\frac{M1-M2}{\sqrt{\frac{{SD1}^{2}+{SD2}^{2}}{2}}}$$

Where *M*1 and *M*2 represent mean values for groups 1 and 2 and *SD*1 and *SD*2 are the corresponding standard deviations. Effect sizes were interpreted small (*d* = 0.2), medium (*d* = 0.5), and large (*d* = 0.8) based on benchmarks suggested by Cohen.

**TEM imaging on endophilin droplets**

For preparing endophilin droplet samples, the protein (25 µM) was incubated with 10% PEG for 30 min. The mixture was allowed to adhere onto formvar coated copper grids (200 mesh) (Electron Microscopy Sciences, Hatfield, PA) for 2 min. Excess liquids were removed from the grids by soaking with Whatmann paper. Grids were stained by 1% (w/v) uranyl acetate solution for 2 min and extra stains were removed by Whatmann paper. Stained grids were washed three times with MiliQ water and air dried for 10 min. Images were recorded on a JEM 1011 transmission EM (JEOL, USA), operated at 100 kV, coupled with an ORIUS 832.10 W CCD camera (Gatan). Post processing of images was performed with ImageJ software.
